# Microbial community structure and microbial networks correspond to nutrient gradients within coastal wetlands of the Laurentian Great Lakes

**DOI:** 10.1101/217919

**Authors:** Dean J. Horton, Kevin R. Theis, Donald G. Uzarski, Deric R. Learman

**Author notes:** Correspondence: Deric R. Learman.

## Abstract

Microbial communities within the soil of Laurentian Great Lakes coastal wetlands drive biogeochemical cycles and provide several other ecosystems services. However, there exists a lack of understanding of how microbial communities respond to nutrient gradients and human activity in these systems. This research sought to address the lack of understanding through exploration of relationships between nutrient gradients, microbial community diversity, and microbial networks. Significant differences in microbial community structure were found among coastal wetlands within the western basin of Lake Erie and all other wetlands studied (three regions within Saginaw Bay and one region in the Beaver Archipelago). These diversity differences coincided with higher nutrient levels within the Lake Erie region. Site-to-site variability also existed within the majority of the regions studied, suggesting site-scale heterogeneity may impact microbial community structure. Several subnetworks of microbial communities and individual community members were related to chemical gradients among wetland regions, revealing several candidate indicator communities and taxa which may be useful for Great Lakes coastal wetland management. This research provides an initial characterization of microbial communities among Great Lakes coastal wetlands and demonstrates that microbial communities could be negatively impacted by anthropogenic activities.

## Introduction

The Laurentian Great Lakes of North America are one of the largest freshwater systems on Earth, and are critical in supporting biogeochemical cycles, freshwater resources, biodiversity, and economic viability of the surrounding region. Notably, the Great Lakes region has been impacted by anthropogenic pressure, with cumulative stress having a particular impact on the western basin of Lake Erie (Danz *et al.,* 2007; Uzarski *et al.,* 2017). These negative impacts extend to ecological transition zones between upland and aquatic environments in the form of coastal wetlands which border the Great Lakes (Uzarski, 2009). Agricultural runoff, atmospheric deposition, and urbanization influence water chemistry, and thereby reduce water quality and impair these coastal wetlands (Trebitz *et al.,* 2007; Morrice *et al.,* 2008). As consequence, research assessing biodiversity and anthropogenic pressure on coastal wetlands of the Great Lakes has surged since the Great Lakes Water Quality Agreement (GLWQA) was established in 1972 (Hackett *et al.,* 2017). While much research on coastal wetlands has flourished in the wake of this international agreement, microbial communities within Great Lakes coastal wetlands remain almost entirely uncharacterized (Hackett *et al.,* 2017). The few research studies on microbial communities in Great Lakes coastal wetlands have focused on the use of microbial enzymatic assays as a tool to explore decomposition rates and nutrient limitation (Jackson *et al.,* 1995; Hill *et al.,* 2006). Community diversity, structure, and taxonomic composition have been largely overlooked. As the microbial communities within Great Lakes coastal wetlands have yet to be fundamentally described, it is important to gather baseline data on what microbes exist within these systems, to elucidate how these microbes could be interacting, and to determine to what extent microbial diversity may already be impacted by anthropogenic chemical disturbance.

Microbial communities contribute substantially to the ecological functioning of coastal wetlands (such as carbon and greenhouse gas cycling, and redox-mediated chemical processes), and these wetlands are vital in the retention of chemical pollutants (e.g., heavy metals), sediments, and excess nutrients (e.g., N and P). Coastal wetlands mitigate the effects of these pollutants and reduce pollution impacts on the Great Lakes themselves (Wang & Mitsch, 1998; Sierszen *et al.,* 2012). Coastal wetlands border much of the Great Lakes coastline, where they make up nearly 200,000 ha of habitat between the United States and Canada, despite reduction of this habitat by approximately 50% since European colonization (Dahl, 1990; Hecnar, 2004; Sierszen *et al.,* 2012). Further, the economy of the Great Lakes is contingent on the existence and proper functioning of coastal wetlands. In providing ecosystem services and promoting biodiversity, these wetlands have an estimated annual worth of $69 billion USD; the value of recreational fishing alone is valued at $7.4 billion USD per year (Krantzberg & de Boer, 2008; Campbell *et al.,* 2015). As such, negative anthropogenic impacts on microbial communities could influence the economic viability of the Great Lakes region, biodiversity retention, and the functioning of critical elemental cycles which commonly occur within freshwater wetlands.

Most notably, carbon mineralization occurs within wetland soils via redox processes mediated by microbial communities, and these processes contribute to pollution mitigation and atmospheric greenhouse gas flux (Conrad, 1996; Reddy & DeLaune, 2008). Wetland soils often become chemically structured with increasing depth through sequential reduction of electron acceptors that decrease in metabolic favorability to microbes due to thermodynamic constraints (Conrad, 1996; Reddy & DeLaune, 2008; Kögel-Knabner *et al.,* 2010). As microbial community metabolism changes in concert with soil chemical profiles, microbial community compositional shifts commonly reflect functional changes of the community (Lüdemann *et al.,* 2000; Edlund *et al.,* 2008; Lipson *et al.,* 2015). However, while availability of electron acceptors may influence chemical and biological structure within wetland soils, concentration of carbon electron donors can influence the vertical stratification of redox processes (Achtnich *et al.,* 1995; Alewell *et al.,* 2008), and by extension, vertical microbial community structure (defined as relative proportions of microbial taxa within a community). As an example of how this may apply to natural environments, increased carbon and nutrient influx from anthropogenic activities (such as agricultural pressure) may impact microbial community structure within coastal wetlands. Impacts to microbial community composition may extend to shifts in chemical cycles and redox processes as consequence, as disturbance to microbial community structure can often lead to a shift in community function (Shade *et al.,* 2012). However, while community structure may be indicative of environmental gradients within wetlands, taxonomic identification of microbes which respond to human pressures is necessary to appreciate which fraction of wetland microbial communities are most sensitive to environmental disturbances.

Networks of microbial taxa exist within microbial communities, and impacts to individual members could affect entire networks (Faust & Raes, 2012). Thus, it is important explore hypothetical microbial networks within natural environments, and their relationships to changing environmental conditions. Understanding how microbial networks respond to physicochemical shifts could aide in predicting how a future change in environmental conditions (perhaps caused by anthropogenic activity) may impact local microbial communities. Further, identifying microbial taxonomic and diversity responses to environmental stressors caused by human activity is the first step in developing biological indicators that can predict levels of anthropogenic stress on natural environments, such as wetlands. Physicochemical and biological indicators have been continuously developed to determine which biological taxa are most sensitive to anthropogenic pressures within freshwater wetlands, and by extension, how these biological responses can inform scientists and managers about the health of coastal wetlands along the Great Lakes (Uzarski *et al.,* 2017). These indices have been established for physical and chemical attributes (such as nutrient levels, urbanization, land use, etc.), as well as several eukaryotic taxonomic groups (e.g., macrophytes, macroinvertebrates, fish, anurans, and birds) (Uzarski *et al.,* 2017). However, as different taxonomic indicators highlight unique pressures on wetland systems, indicators based on different biological groups can often conflict in their assessment of wetland ecosystem health. As such, it is necessary to examine a wide range of biological indicators to assess different aspects of wetland ecosystem health. A biological index for bacteria and archaea has yet to be developed for responses to human impacts within freshwater coastal wetlands (Uzarski et al., 2017). A first step in establishing a microbial index is to uncover specific networks of microbial taxa (Sims *et al.,* 2013; Urakawa & Bernhard, 2017) and diversity patterns found to be related to environmental gradients linked to anthropogenic activity (e.g., soil nutrient levels) among Great Lakes coastal wetlands.

This study sought to provide an initial characterization of microbial communities within soils of Great Lakes coastal wetlands bordering the western basin of Lake Erie, Saginaw Bay of Lake Huron, and northern Lake Michigan. Wetland sites explored in this study have been extensively researched over multiple years and vary widely in the degree to which they are impacted by human activity (Uzarski *et al.,* 2017). This study explored how environmental gradients among these coastal wetlands were related to microbial community structure among wetlands. Additionally, relationships among microbial communities and changing environmental conditions with increasing soil depth were also explored within each wetland site. It was predicted that microbial community structure would be related to environmental gradients among and within coastal wetland regions of the Great Lakes, and elevated nutrient levels within wetlands would decouple the relationship between microbial community structure and soil depth with respect to coastal wetlands lower in nutrient levels, as has been suggested in previous studies (Achtnich *et al.,* 1995; Alewell *et al.,* 2008). Through high-throughput sequencing of the 16S rRNA gene and microbial network analyses, variations in key microbial taxa and subcommunities related to environmental gradients established by wetlands were identified.

## Materials and Methods

### Study site and field sampling

In the summer of 2014, wetland soil cores were collected within Laurentian Great Lakes coastal wetland ecosystems. Specifically, soil cores were collected from ten sites across five regions, including two sites in the western basin of Lake Erie (LE), three sites in eastern Saginaw Bay (ESBT), two sites in northern Saginaw Bay (NSB), two sites in western Saginaw Bay (WSB) in Lake Huron, and one site in the Beaver Island archipelago (BA) in Lake Michigan (Fig. 1). These sites were selected as they corresponded to environmental gradients, as well as human impact gradients based upon SumRank scores (an index assessing land use and water quality) as described in Uzarski *et al.* (2017) (Supplemental Fig. 1). Soil cores were collected by hand-driving plastic core tubes (~ 5 cm diameter) vertically into the soil. Among wetlands, samples were collected within the same vegetation zone across sites (either dominated by cattails, genus *Typhus,* or bulrush, genus *Shoenoplectus*) as an attempt to control for collection bias, as different vegetation zones can harbor microbial communities distinct from other vegetation zones (Tang *et al.,* 2011). Cores were sampled to a depth of at least 6 cm (except for one core which was sampled to a depth of 4 cm) and were immediately flash frozen in a dry ice ethanol bath. Samples were transported on dry ice to Central Michigan University wherein they were stored at −80 °C.

**Figure 1.**
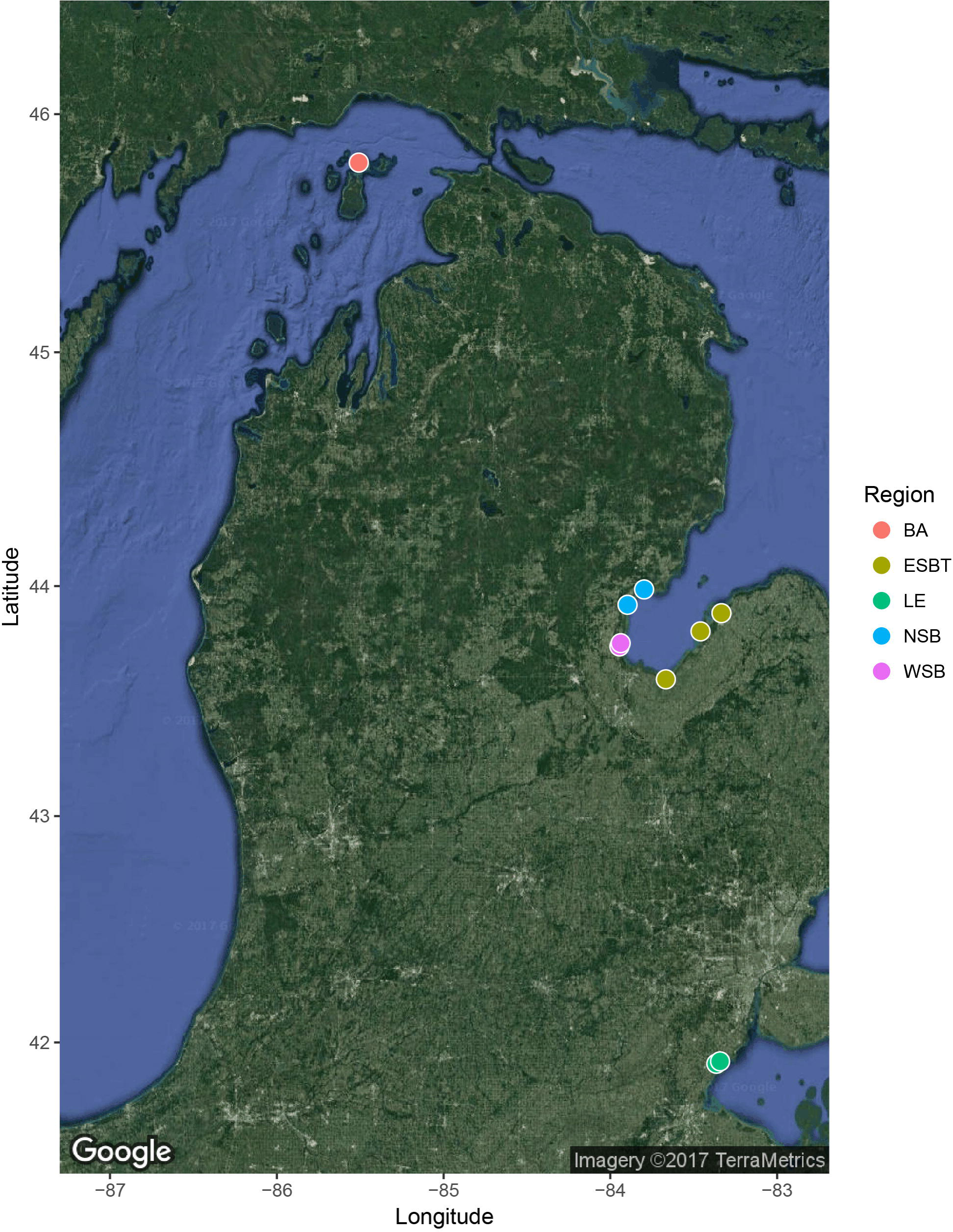
Geographic map displaying locations of sites sampled within this study. Colors of points correspond to region sampled.

Triplicate cores were taken at five wetland sites while duplicate cores were taken at five other wetland sites. Global Positioning System (GPS) coordinates were recorded at each sampling location. For sample extraction and sectioning, cores were extruded while still frozen via a custom-built core extruder. The edge of the core was warmed with a heat gun to allow the soil core to pass efficiently through the plastic container, however, the inner-core did not thaw during extrusion. Ice was applied to the plastic core liner to prevent accelerated thawing. Beginning from the top surface of soil, 2 cm sections were cut via an ethanol and flame-sterilized hacksaw blade and the sectioned core samples were placed into Whirl-Pak bags and stored at −80 °C. The extruder face plate was sterilized between cuts of the same core with ethanol. The extruder device was fully cleaned and sterilized between cores with physical scrubbing and ethanol sterilization.

### Microbial community analysis

Each soil sample was analyzed independently for microbial community analyses. DNA was extracted from ~ 0.25 g of soil using a MoBio PowerSoil DNA Isolation Kit (Mo Bio, Carlsbad, CA) following the standard manufacturer’s protocol. Concentrations of extracted DNA were assessed using a Qubit^®^ 2.0 fluorometer (Life Technologies, Carlsbad, CA) to ensure successful DNA extraction and quantification for sequence library preparation. DNA samples were sent to Michigan State University (East Lansing, MI) for library preparation and sequence analysis at the Research Technology Support Facility. The V4 region of the 16S rRNA gene was amplified for downstream sequencing with the commonly used primers 16Sf-V4 (515f) and 16Sr-V4 (806r) and a previously developed protocol (Caporaso *et al.,* 2012; Kozich *et al.,* 2013). Paired-end 250 bp sequencing was accomplished via a MiSeq high-throughput sequencer (Illumina, San Diego, CA). Acquired DNA sequences were filtered for quality and analyzed using MOTHUR v 1.35.1 (Schloss *et al.,* 2009) following the MiSeq SOP (available at https://www.mothur.org/) with modifications. Scripts used for sequence processing can be found at the GitHub repository associated with this study (https://github.com/horto2dj/GLCW/). Briefly, paired end sequences were combined into single contigs. Sequences that contained homopolymers > 8 bases, and those less than 251 or greater than 254 bp were removed. Sequences were aligned against the Silva (v 119) rRNA gene reference database (Quast *et al.,* 2012). Sequences which did not align with the V4 region were also subsequently removed from analysis. Chimeric DNA was searched for and removed via UCHIME (Edgar *et al.,* 2011). Sequences were classified via the Ribosomal Database Project (training set v 9; Cole *et al,* 2013) with a confidence threshold of 80. Sequences classified as chloroplast, mitochondria, eukaryotic, or unknown were removed. Remaining sequences were clustered into Operational Taxonomic Units (OTUs) at 0.03 sequence dissimilarity using the opticlust clustering algorithm. Sequence data associated with this research have been submitted to the GenBank database under accession numbers SRR6261304 – SRR6261377 (Horton et al., 2017).

### Chemical analysis

Each soil layer (top, middle, and bottom) was analyzed separately for local chemistry at each site. Within each site, soil samples of the same depth (i.e., top, middle, and bottom soil samples) among duplicate/triplicate cores were combined and homogenized to obtain enough soil for chemical analyses. For chemical analysis, soil samples were sent to Michigan State University Soil & Plant Nutrient Lab (East Lansing, MI) to analyze for percent total N (“TN”), total P (“TP”, ppm), total S (“TS”, ppm), NO_3_^-^ (ppm), NH_4_+(ppm), percent organic matter (“OM”), percent organic carbon (“OC”), and C:N. In the field, a YSI multiprobe (YSI Inc., Yellow Springs, OH) was used to measure pH of the water residing directly above each collected soil core. Other data generated for this study, along with R code for replication of statistical methodology, can be found in the GitHub repository at https://github.com/horto2dj/GLCW/.

### Statistical analyses

Statistical analyses were completed using R statistical software version 3.2.2 (R Core Team, 2015) unless otherwise stated. Code used for statistical analyses (and bioinformatic workflow) in this study can be found in the associated GitHub repository (https://github.com/horto2dj/GLCW/).

#### Physicochemical analysis

Differences in chemical profiles between samples within and among wetland regions were visualized using Principal Component Analysis (PCA). Prior to PCA, percentages were arcsin square root transformed and ratios were log transformed. Additionally, Pearson correlation analyses were performed to search for significant correlations between chemical variables. Collinearity in the dataset was addressed by combining highly correlated environmental variables (r > 0.7, p ≤ 0.001). Only one of the correlated variables was included in PCA to remove exaggeration of correlated variables in PCA structure. Permutational Multivariate Analysis of Variance (perMANOVA; Anderson, 2001) was used to determine the influences of region and soil depth on physicochemical composition of samples, and 95% confidence intervals were established to compare differences among groups. Chemical depth profiles were also visualized for each wetland site to understand shifts in measured environmental variables with increasing soil depth.

#### Alpha diversity analysis

Alpha diversity analyses were performed to explore variation in OTU richness and evenness among wetland sites, regions, and soil depths, as well as to determine whether observed trends were driven by environmental variables. Prior to alpha diversity analyses, sequence abundance for each sample was subsampled to the lowest sequence abundance for any one sample (n = 48,226 sequences). Singletons were maintained within the sequence dataset for alpha diversity analyses, as alpha diversity indices can be reliant on the presence of singletons for proper estimation. Alpha diversity was calculated for each site using MOTHUR, including Chao1 richness and non-parametric Shannon diversity. Linear mixed-effect models and ANOVAs were used to test influences of wetland site, region, and soil depth on alpha diversity, controlling for wetland site as a random effect. Linear models and ANOVAs were used to test for variation in alpha diversity among wetland sites. If significant variation was found within an ANOVA result, post-hoc comparisons were implemented between sample groups using Tukey’s Honest Significant Differences (HSD) tests with Bonferroni adjustments (p-values obtained by number of comparisons) for pairwise comparisons.

#### Beta diversity analysis

Beta diversity analyses were used to evaluate variation in microbial community structure among wetland sites, regions, and soil depths, and to assess the extent to which observed variation was explained by environmental conditions. Singletons and doubletons were removed from the dataset for beta diversity analyses. All sequence data were maintained for beta diversity analyses and transformed using the *DeSeq2* (Love *et al.,* 2014) package, which normalized OTU abundances among samples using a variance stabilizing transformation (VST) (McMurdie & Holmes, 2014). The *phyloseq* (McMurdie & Holmes, 2013) and *Vegan* (Oksanen *et al.,* 2007) packages were used to compare beta diversity among samples. Dissimilarity in microbial community structure among samples within and among sites was visualized using Non-metric Multidimensional Scaling (NMDS) plots based on pairwise Bray-Curtis dissimilarity estimates. The function *envfit* of the Vegan package was used to evaluate correlation between chemical parameters and microbial community structure among samples according to NMDS. “Depth” was also implemented as a dummy variable to test correlation between depth and microbial community structure.

To test for significant differences in beta diversity among wetland sites, regions, and soil depth, perMANOVA were implemented. Specifically, these tests evaluate significant variation among within group and between group means (Clarke, 1993; Anderson, 2001; Anderson & Walsh, 2013). If perMANOVA found significant differences among groups at the global level, pairwise perMANOVA tests between groups were implemented with Bonferroni significance adjustments to control for multiple pairwise comparisons. Anderson’s permutation of dispersions test (PERMDISP; Anderson, 2006; Anderson *et al.,* 2006) was used to test for differences in variance of community structure among sample groups (i.e. sites, regions, soil depths). Tukey’s Honest Significant Difference (HSD) tests were implemented with adjusted p-values for multiple pairwise comparisons if significant differences in dispersion were found among groups.

To explore relationships between regional microbial community structure and environmental variables, NMDS plots were generated for each individual region. Applying NMDS to each region also allowed for the assessment of the correlational relationship between community structure and soil depth (as a dummy variable) and other environmental variables (using the *envfit* function) within individual regions. To test for differences in microbial community structure between/among sites within a region, as well as among depths within a region, perMANOVA was implemented individually for each region.

#### Taxonomic analyses

Dominant microbial taxa were explored in order to characterize microbial communities within Great Lakes coastal wetlands. Differential abundance analysis was performed for microbial OTUs between significantly different wetland regions and soil depths (according to perMANOVA results with all microbial samples included) using the *DESeq2* package. OTUs which did not appear at least twice within 10% of samples explored and were not significantly differentially abundant at p < 0.001 were omitted from differential analyses to minimize spurious relationships.

#### Network analyses

To explore relationships between microbial sub-communities and individual OTUs to environmental variables, Weighted Correlation Network Analysis (WGCNA) was implemented on OTU relative abundances using the *WGCNA* package (Langfelder & Horvath, 2008; Langfelder & Horvath, 2012), executed as previously described (Guidi *et al.,* 2016; Henson *et al,* 2018) with modifications. OTUs which did not possess at least 2 sequences across 10% of samples were removed from network analyses. These OTUs were removed to eliminate OTUs with potentially spurious correlations to environmental variables or other OTUs, as well as to reduce computational stress of analyses. Remaining OTU abundances across samples were normalized using variance stabilizing transformation (VST) performed as described previously for beta diversity analyses. To ensure scale-free topology of the network, the dissimilarity matrix generated through VST was transformed to an adjacency matrix by raising this dissimilarity matrix to a soft threshold power. A threshold power of *p* = 4 was chosen to meet scale-free topology assumptions based upon criterion established by Zhang & Horvath (2005). Scale-free topology of network relationships was further ensured through regression of the frequency distribution of node connectivity against node connectivity; a network is scale-free if an approximate linear fit of this regression is evident (see Zhang & Horvath, [2005] for more indepth explanation). A topological overlap matrix (TOM) was generated using the adjacency matrix, and subnetworks of highly connected and correlated OTUs were delineated with the TOM and hierarchical clustering. Representative eigenvalues of each subnetwork (i.e., the first principal component) were correlated (Pearson) with values of measured environmental variables to identify the subnetworks most related to said environmental variables. The subnetworks with the highest positive correlations to environmental variables of interest (e.g., NO_3_^-^, C:N, etc.) were selected for further analyses of relationships among subnetwork structure, individual OTUs, and environmental variables. Partial Least Square regression (PLS) was used to test predictive ability of subnetworks in estimating variability of environmental parameters, which allowed for delineation of potential indicator subnetworks and OTUs. Pearson correlations were calculated between response variables and leave-one-out cross-validation (LOOCV) predicted values. If PLS found that regression between actual and predicted values was below minimum threshold of *R^2^* = 0.3, WGCNA analysis was halted for that network, as the network was deemed to lack predictive ability of that environmental variable. Variable Importance in Projection (VIP) (Chong & Jun, 2005) analysis was used to determine the influence of individual OTUs in PLS. A high VIP value for an OTU indicates high importance in prediction of the environmental response variable for that OTU. For network construction and visualization purposes, the minimum correlation value required between two OTUs to constitute an “edge” between them was delineated at different *r* values for each network related to an individual environmental variable (ranging between 0.1 – 0.25), as co-correlations between OTUs within some networks were stronger than others. The number of co-correlations an OTU has with other OTUs within a network defines its “node centrality” (as described by Henson *et al.,* 2018).

## Results

### Chemical analyses

Significant correlations (r > 0.7, p ≤ 0.001) were found among NH_4_^+^, OM, OC, TN, and latitude. Thus, downstream analyses combined these values into one parameter, “NUTR”, represented by OM values as this variable was the most strongly correlated with each of the other variables. Environmental data were analyzed with a PCA and PC1 and PC2 explained 56.2% and 20.6% of the variation among samples, respectively (Fig. 2). perMANOVA found significant differences in physicochemical profiles based on region (R^2^ = 0.570, p ≤0.001) and depth (R^2^ = 0.058, p ≤ 0.01). Lake Erie coastal wetlands were chemically distinct from other wetland regions (ESBT and NSB; adjusted p = 0.01) according to perMANOVA and pairwise perMANOVA based on Euclidean distance. Ninety-five percent confidence intervals demonstrated no overlap between Lake Erie coastal wetlands and other coastal wetlands (Fig. 2). This separation was related to increased NUTR, NO_3_^-^, and S.

**Figure 2.**
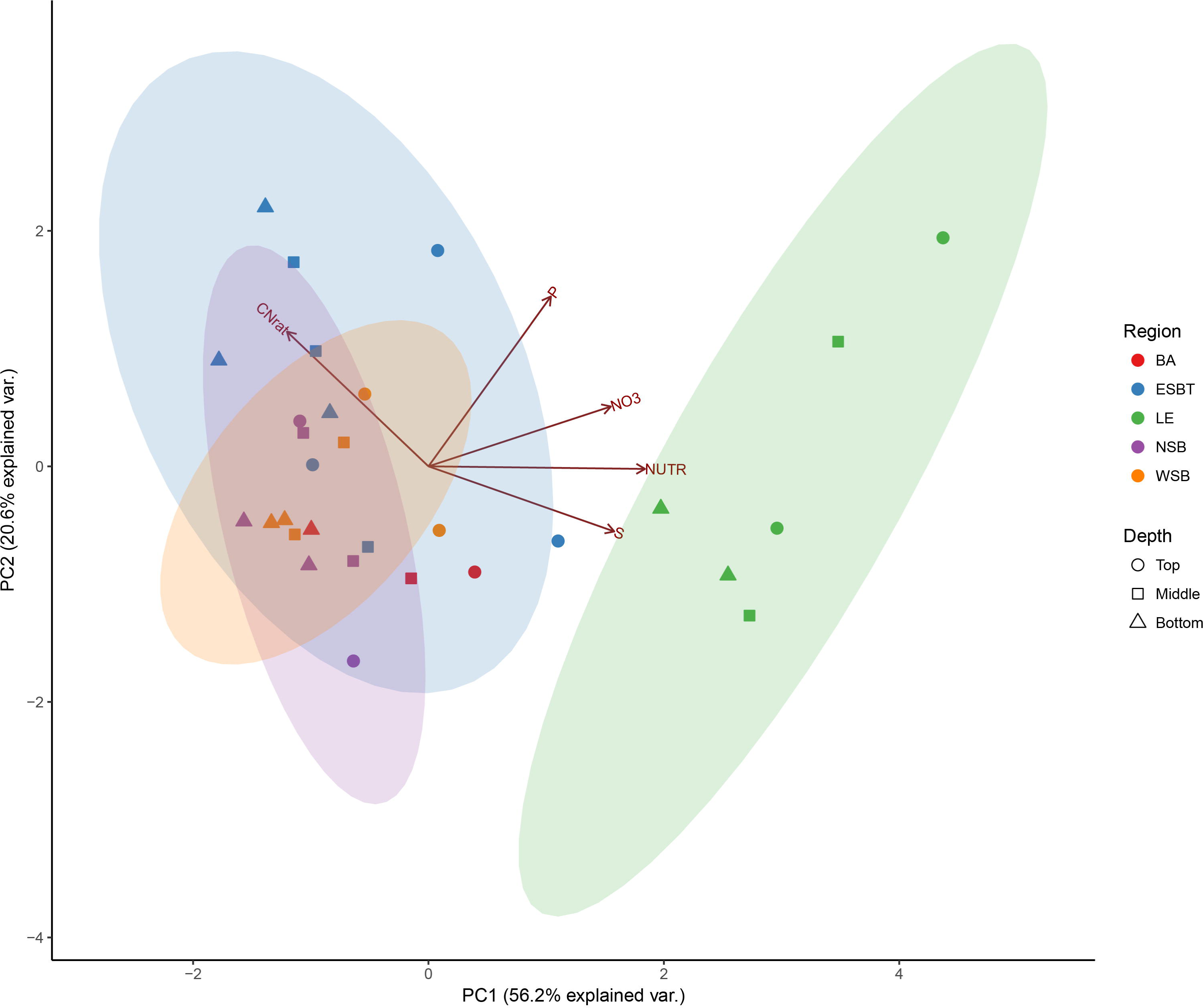
Principal Component Analysis (PCA) illustrating separation of samples based upon soil geochemistry. Shapes and colors correspond to different wetland depths and regions, respectively, as listed in the legend. Percentages on axes represent explained variance of that principal component. Vectors represent impact of specific environmental variables on sample distribution. NUTR represents OM values, which correlated significantly (p ≤ .01, r > 0.56) to NO_3_^-^, OC, OM, S, and TN. Ellipses represent 95% confidence intervals of regional groupings.

Increasing depth within cores showed a consistent shift in environmental variables, except in those sites located in the western basin of Lake Erie (Supplemental Fig. 2). Specifically, OM, OC, and TN consistently decreased with increasing depth within each region except Lake Erie. Similarly, C:N increased with depth in each region except Lake Erie, wherein the C:N ratio remained relatively low (~ 12) and stable with increasing soil depth. Within the Lake Erie wetland region, pH was more acidic in the overlying water with respect to all other wetland regions (Supplemental Table 1). However, pH was still relatively neutral within Lake Erie (average pH = 7.26 ± 0.24), whereas other wetland regions (regions within Saginaw Bay and Beaver Archipelago) experienced slightly more basic pH, with average pH among these regions ranging between 7.72 – 8.39.

### Alpha diversity

Sufficient depth of sampling was reinforced by rarefaction curve analysis (Supplemental Fig. 3). Good’s coverage values ranged between 89.3 – 93.5% for each region at the subsampled value of 48,226 sequences. Chao1 richness estimates varied significantly among wetland regions (F = 8.38, p ≤ 0.05), as well as wetland sites (F = 16.78, p ≤ 0.001). Pairwise comparisons found that the LE region had significantly higher (p ≤ 0.01) Chao1 estimates than NSB and WSB regions (Fig. 3; Supplemental Table 2). Additionally, pairwise comparisons found a high degree of significant variability (p ≤ 0.01) in Chao1 estimates among wetland sites (Supplemental Table 2). Further, Shannon diversity levels also significantly varied among wetland sites (F = 4.57, p ≤ 0.001), with site LE_D having significantly higher (p ≤ 0.01) Shannon diversity levels than sites ESBT_A and WSB_B (Supplemental Table 2). Soil depth did not influence alpha diversity levels.

**Figure 3.**
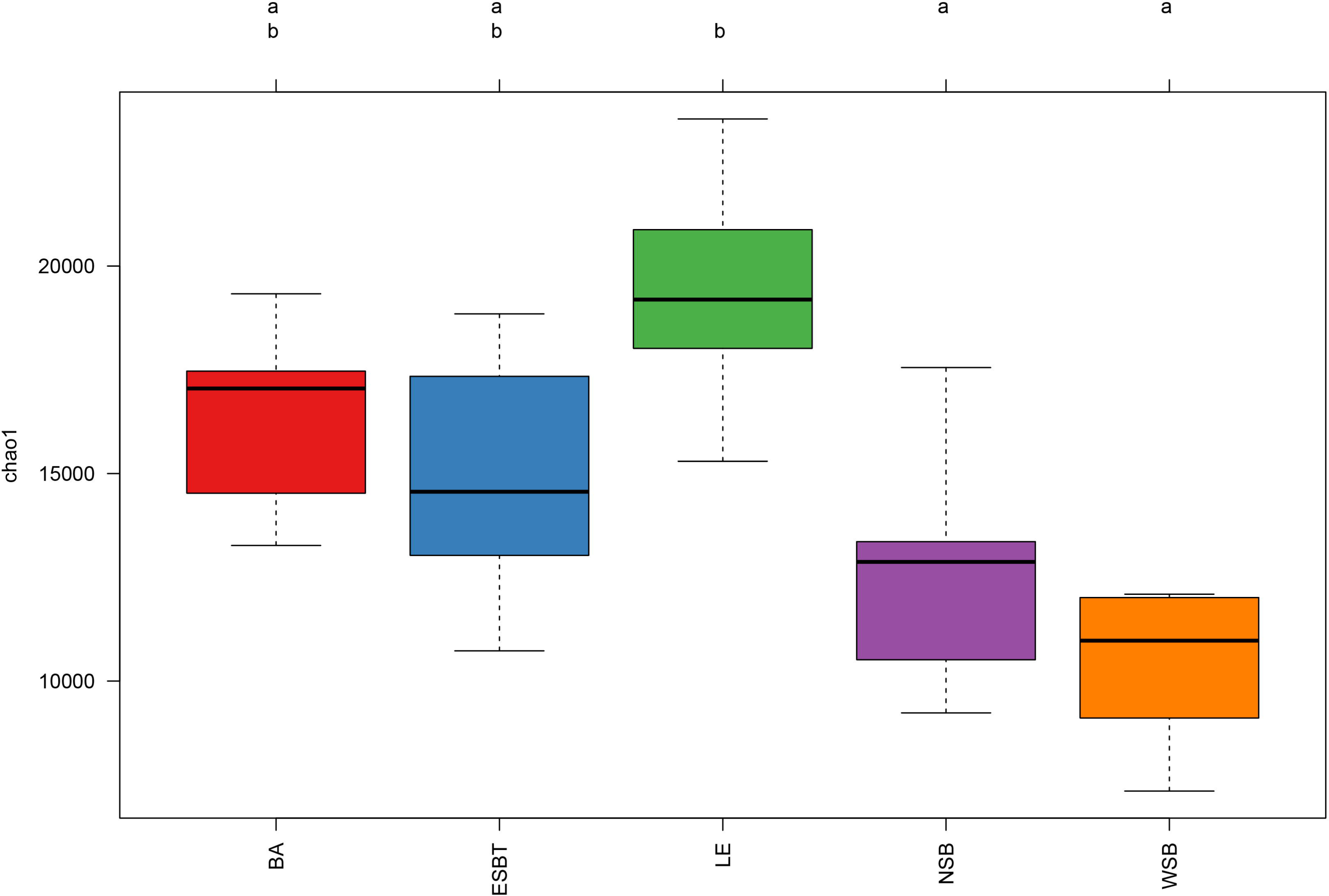
Boxplot diagram comparing Chao1 diversity among wetland regions. Boxes with the same letter are not significantly different, while those with no common letters are significantly different (p ≤ 0.01). Lines within boxes represent the median, hinges represent +/- 25% quartiles, whiskers represent up to 1.5x the interquartile range. Colors represent wetland region.

Shannon diversity and Chao1 were both positively correlated with measured environmental variables (Table 1). Specifically, Chao1 estimates increased with NO_3_^-^, P, and S concentrations (p ≤ 0.01), and were weakly positively correlated (p ≤ 0.05) with NUTR. Additionally, Shannon diversity levels increased alongside NUTR and S (p ≤ 0.001), and were weakly positively correlated with NO_3_^-^ (p ≤ 0.05). There were no significant relationships between alpha diversity and C:N, and alpha diversity was not negatively correlated with any of the measured environmental variables.

**Table 1.** Correlations between alpha diversity metrics and measured environmental variables. Asterisks represent significance values where p < 0.001 (***), p < 0.01 (**), and p < 0.05 (*).

|  | Chao1 | Shannon |
| --- | --- | --- |
| P | 0.31** | -0.09 |
| S | 0.42*** | 0.45*** |
| NO<sub>3</sub><sup>-</sup> | 0.42*** | 0.24* |
| C:N | -0.03 | -0.2 |
| NUTR | 0.24* | 0.41*** |

### Beta diversity

#### Beta diversity among regions

Multivariate analyses were implemented to explore relationships between microbial communities and environmental gradients among wetland regions. NMDS demonstrated separation of microbial communities based on wetland site, region, and soil depth (Fig. 4). Substantiating this result, perMANOVA confirmed that differences in microbial community structure were significantly related to wetland region (R^2^ = 0.220, p ≤ 0.001), site (R^2^ = 0.119, p ≤ 0.001), and soil depth (R^2^ = 0.070; p ≤ 0.001). Post-hoc pairwise perMANOVA found that community structure within the LE region was significantly distinct (p ≤ 0.01) from all other wetland regions (Table 2). No significant differences in community structure were found between any other wetland regions compared. Additionally, microbial community beta diversity was distinct (p ≤ 0.003) between the top soil depth and the middle and bottom soil depths. However, no significant differences in microbial community structure were found between the middle and bottom soil depths (Table 2). Variation in microbial community structure was significantly correlated (p ≤ 0.001) to depth (r = 0.41), NO_3_^-^ (r = 0.20), NUTR (r = 0.60), and S (r = 0.41), and also correlated (p ≤ 0.016) with C:N (r = 0.11) and P (r = 0.14) (Supplemental Table 3).

**Figure 4.**
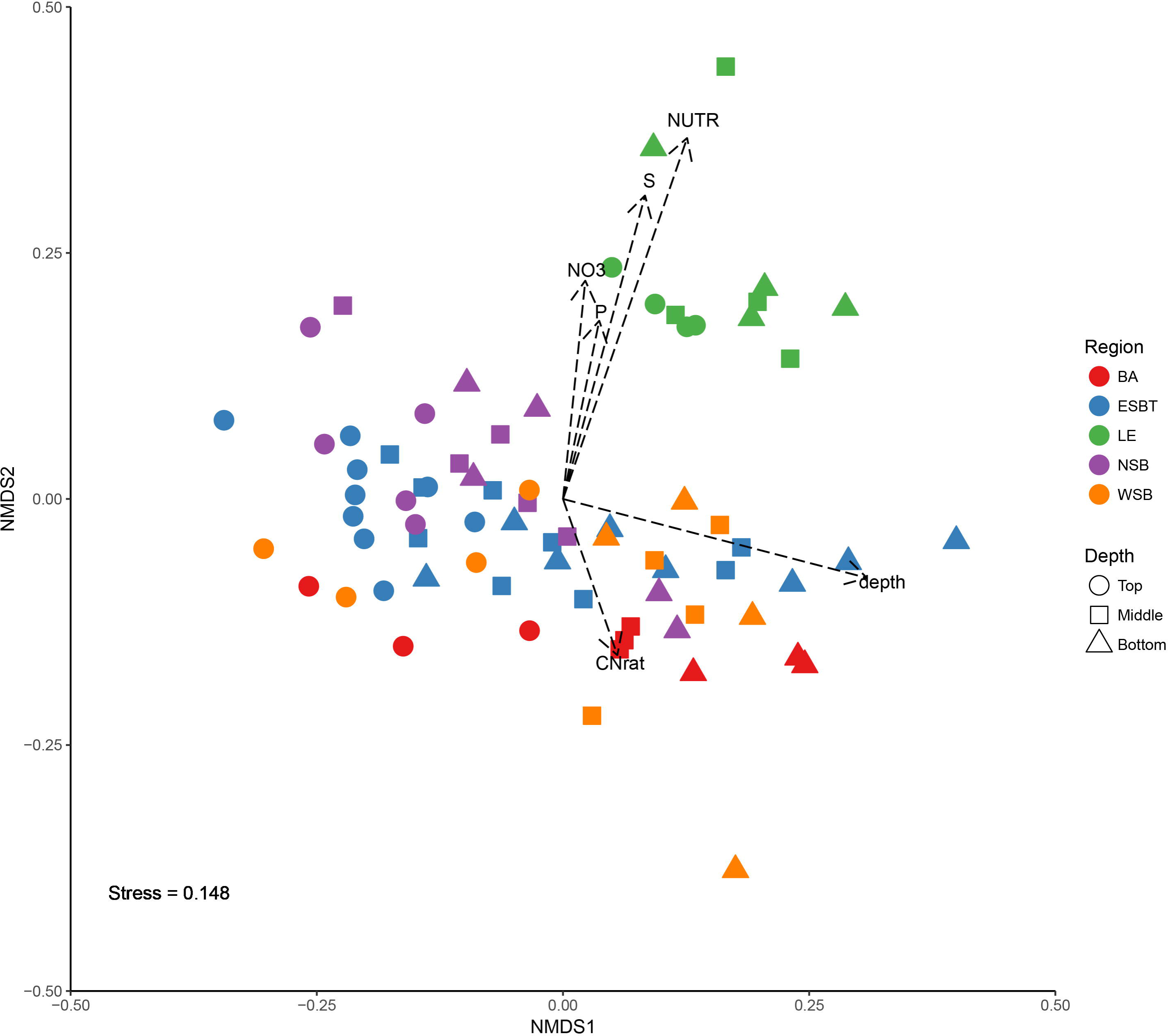
Nonmetric Multidimensional Scaling (NMDS) plot illustrating separation of samples based upon differences in microbial community structure. Shapes and colors correspond to different depths and wetland regions, respectively, as listed in the legend. Vectors represent correlations of environmental variables to the distribution of the microbial communities represented in the plot.

**Table 2.** Pairwise perMANOVA results comparing pairwise differences between wetland regions and differences between wetland soil depths. Values represent significant (p < 0.01) R^2^ results, and *n.s.* represents lack of significance (p > 0.01).

| Region | BA | ESBT | LE | NSB | WSB |
| --- | --- | --- | --- | --- | --- |
| BA | - |  |  |  |  |
| ESBT | *n.s.* | - |  |  |  |
| LE | 0.507 | 0.401 | - |  |  |
| NSB | *n.s.* | *n.s.* | 0.524 | - |  |
| WSB | *n.s.* | *n.s.* | 0.435 | *n.s.* | - |
| Depth | Top | Middle | Bottom |  |  |
| Top | - |  |  |  |  |
| Middle | ** | - |  |  |  |
| Bottom | ** | *n.s.* | - |  |  |

Beta dispersion tests suggested significant variation in structural variance among regions (p ≤ 0.05), however, Tukey’s HSD test using adjusted p-values for multiple comparisons did not find any significance (p > 0.05) between pairwise comparisons of regional groups. There were no differences in community structural dispersion among soil depths.

#### Beta diversity within regions

Microbial community associations with environmental variables were also explored within regions to examine variation among wetland sites. Individual NMDS plots of each region identified relationships between microbial community structure and several environmental variables using vector-fitting regression, and strengths of these relationships were dependent upon the wetland region explored (Fig. 5; Supplemental Table 3). Depth was significantly related (p ≤ 0.05) to microbial community structure in all wetland regions except NSB and LE. However, microbial community structure may have been more strongly related to depth in NSB (r = 0.35, p = 0.071) than LE (r = 0.19, p = 0.40). NUTR was significantly related (p ≤ 0.01) to community structure within regions BA (r = 0.82), ESBT (r = 0.51), and LE (r = 0.66). C:N was related (p ≤ 0.01) to community structure within regions of Saginaw Bay (i.e., ESBT [r = 0.65], NSB [r = 0.58], and WSB [r = 0.58]). Beta diversity was not significantly associated with concentrations of NO_3_^-^ in any region.

**Figure 5.**
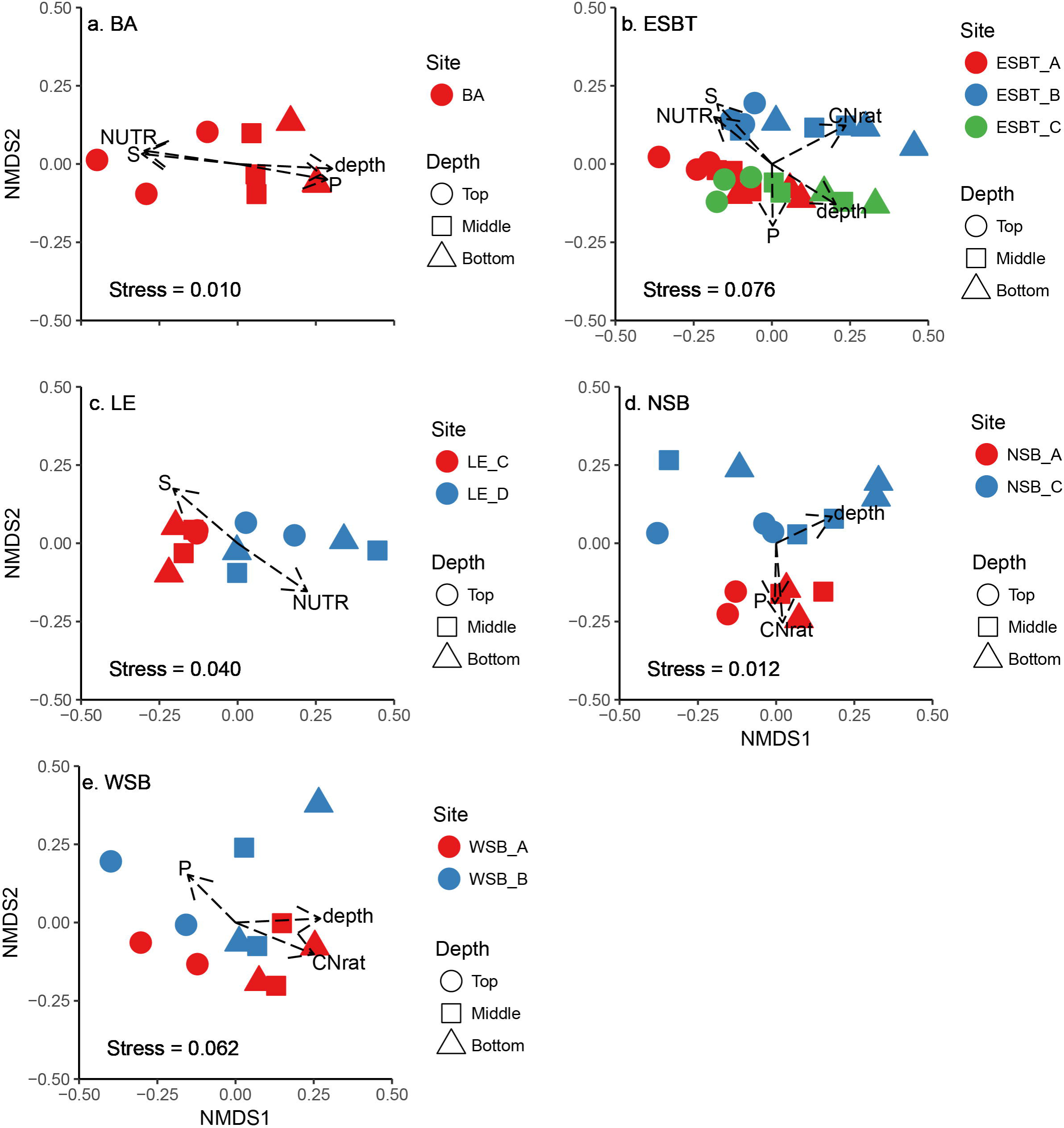
NMDS plots of each wetland region demonstrating separation of samples based upon differences in microbial community structure, including **(A)** BA, **(B)** ESBT, **(C)** LE, **(D)** NSB, and (E) WSB. Shapes and colors correspond to different depths and wetland sites, respectively, as listed in the legends. Vectors represent correlations of environmental variables to the distribution of microbial communities represented in the plots.

To test for significant differences in microbial beta diversity within regions, perMANOVA was used to evaluate differences in microbial community structure among soil depths and sites within wetland regions (Supplemental Table 3). Depth did not significantly explain microbial community structure within the region LE (p = 0.65), however, it did explain differences in microbial community structure within the other wetland regions, specifically BA (R^2^ = 0.414; p = 0.006), ESBT (R^2^ = 0.154; p = 0.001), NSB (R^2^ = 0.161; p = 0.093), and WSB (R^2^ = 0.259; p = 0.014). Significant differences in microbial community structure were found among different wetland sites within regions ESBT (R^2^ = 0.192; p = 0.001), LE (R^2^ = 0.236; p = 0.004), and NSB (R^2^ = 0.140; p = 0.003). As only one site was sampled within the BA region, testing for differences among wetland sites within the BA region could not be accomplished.

### Taxonomic analyses

At the level of Phylum, wetland sites were dominated by similar consortia of bacteria and archaea. Soils had a high relative abundance of *Proteobacteria,* with *Deltaproteobacteria* and *Betaproteobacteria* comprising the largest fraction of *Proteobacteria* (ranging between 7 – 15% of tall taxa; Supplemental Fig. 4). Other relatively abundant bacteria included the phyla *Bacteroidetes, Chloroflexi, Verrucomicrobia, Firmicutes, Acidobacteria, Chlorobi, Actinobacteria,* and *Planctomycetes,* and the classes *Gammaproteobacteria* and *Alphaproteobacteria* within the phylum *Proteobacteria.* One archaeal phyla, *Euryarchaea,* was abundant within wetland soils, ranging between 2 – 5% relative abundance within each wetland site. Between 21 – 32% of bacterial and archaeal taxa among sites were unclassified.

Differential analysis comparing the LE region to all other wetland regions (i.e., BA, ESBT, NSB, and NSB) identified 1,182 OTUs which were differentially abundant across 44 Classes within 15 Phyla (Fig. 6). Differential analysis comparing the top section of wetland soil to the bottom section of wetland soil found 516 OTUs which were differentially abundant between the two zones across 33 Classes within 15 Phyla (Fig. 7).

**Figure 6.**
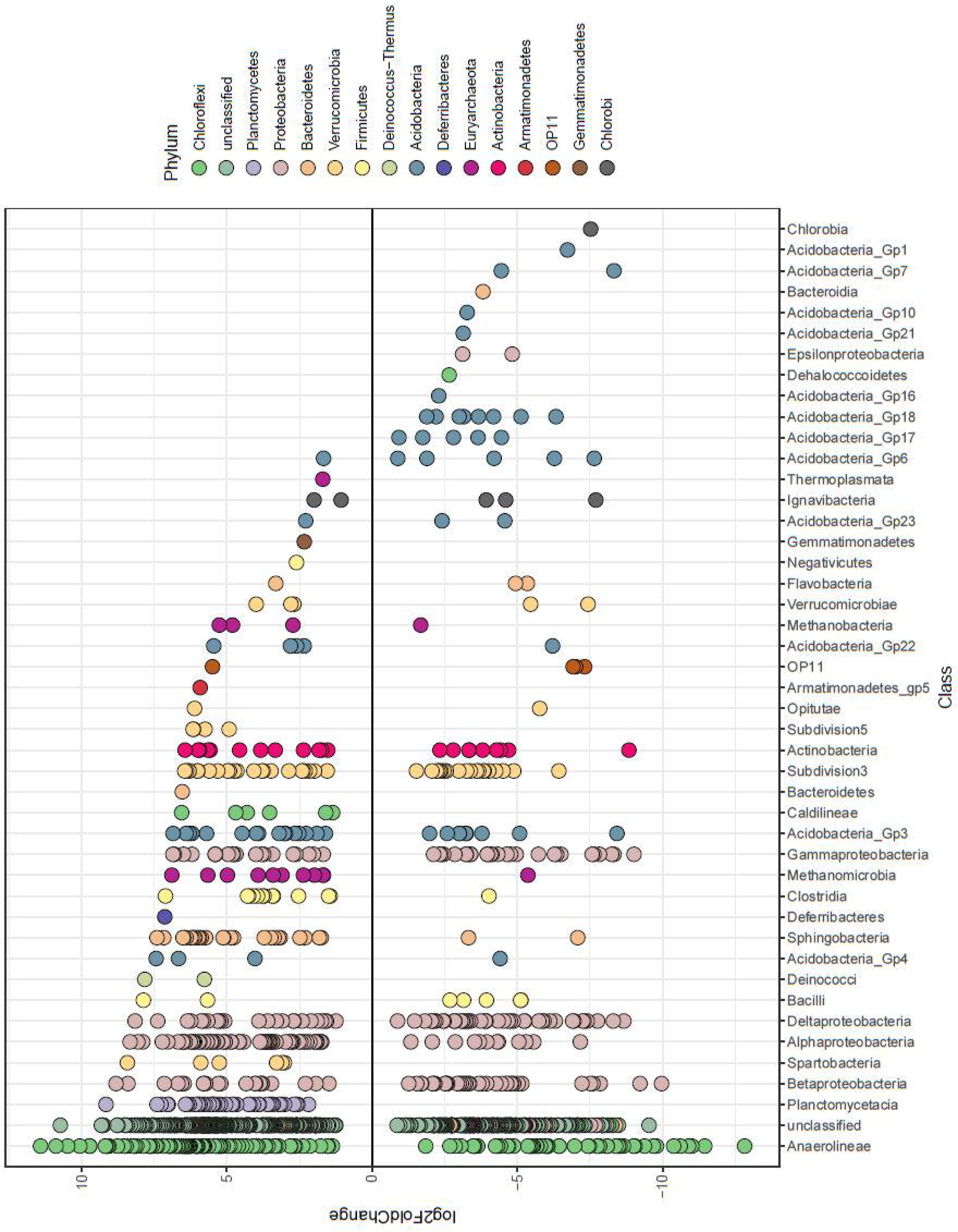
Differential analysis results comparing differentially abundant OTUs between the LE region and all other wetland regions (i.e., BA, ESBT, NSB, WSB). Points represent individual OTUs, and OTU placement above or below the “0” line represents an OTU’s corresponding logarithmic fold change at log_2_. OTUs below the “0” line represent OTUs which were more relatively abundant within the LE region, and OTUs above the “0” line represent OTUs which were more relatively abundant within other wetland regions. Color of point represents phylum identity, and columns represent the Class to which an OTU was confidently assigned (bootstrap value of 100).

**Figure 7.**
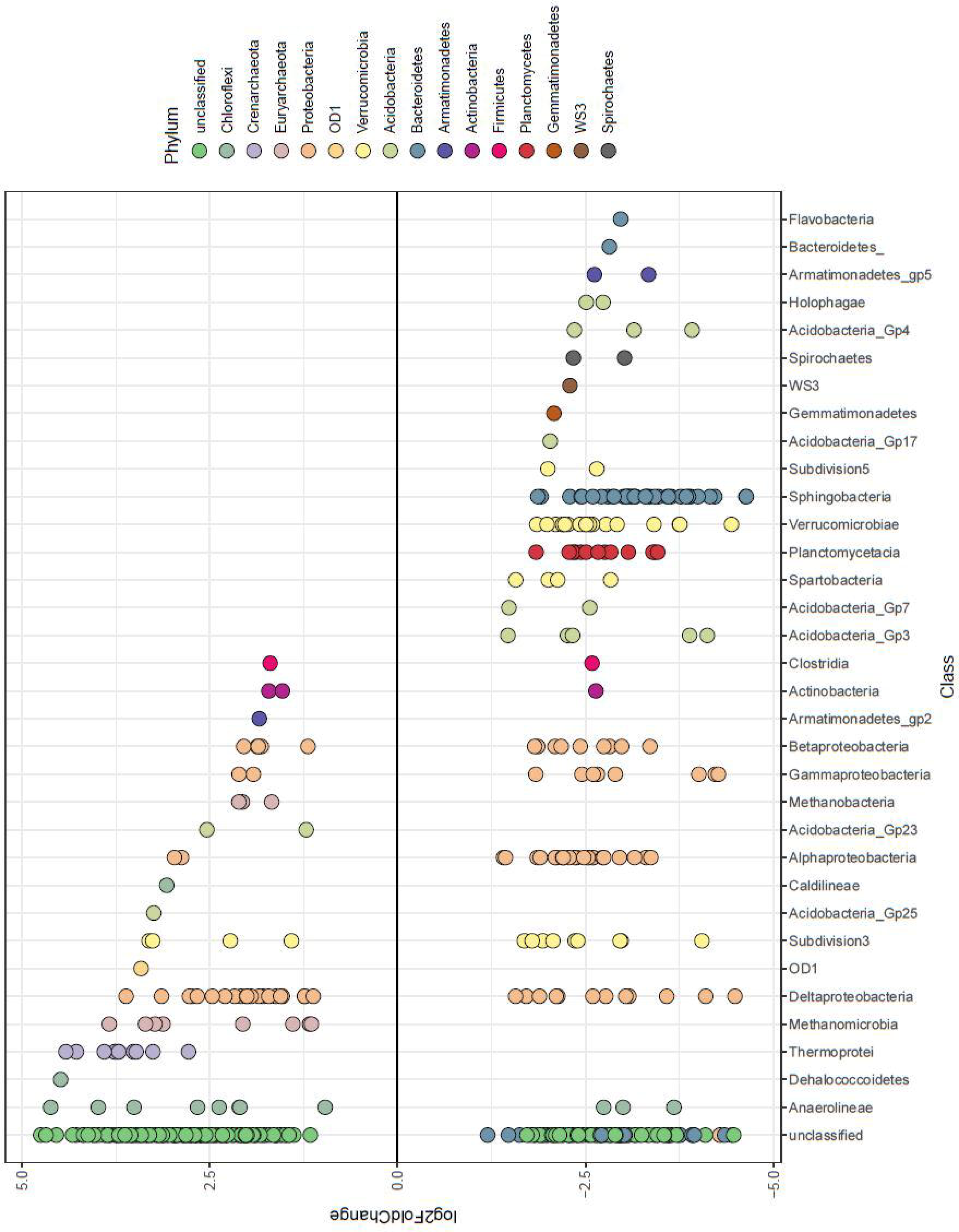
Differential analysis results comparing differentially abundant OTUs between the top and bottom wetland soil zones. Points represent individual OTUs, and OTU placement above or below the “0” line represents an OTU’s corresponding logarithmic fold change at log_2_. OTUs below the “0” line represent OTUs which were more relatively abundant within the top soil layer (0 – 2 cm), and OTUs above the “0” line represent OTUs which were more relatively abundant within the bottom soil layer (4 – 6 cm). Color of point represents phylum identity, and columns represent the Class to which an OTU was confidently assigned (bootstrap value of 100).

### Network analyses

Weighted Correlation Network Analysis (WGCNA) was used to explore strong relationships between subcommunities and individual OTUs with environmental parameters within Great Lakes coastal wetlands. After removal of OTUs that did not have at least two representative sequences in at least 10% of samples, a total of 7,562 OTUs remained for WGCNA. In determining scale-free topology of the OTU network, a soft power threshold of 4 was reached, and an R^2^ of 0.87 was established as linear fit from the regression of the frequency distribution of node connectivity against node connectivity (Supplemental Fig. 5). Of the 33 constructed subnetworks, the same one (subnetwork “orange”) was found to be most strongly correlated to both NUTR (r = 0.94) and NO_3_^-^ (r = 0.55) (Supplemental Fig. 6). A separate subnetwork (“pink”) was strongly correlated (r = 0.74) to C:N. All correlations of subnetworks to environmental variables were significant (p ≤ 0.001). OTU VIP values ≤ 1 were removed due to the large amount of OTUs within subnetworks correlated with C:N for visualization purposes.

For subnetwork relationships to NUTR (including OM, OC, NH_4_+, and TN), partial least squares analysis (PLS) found that 69 OTUs were 93.8% predictive of variance in NUTR (Supplemental Fig. 7). OTU co-correlation networks were constructed using an OTU co-correlation threshold of 0.25, with strong correlations (r > 0.59) between all OTUs and NUTR (Fig. 8). Of the top 15 OTUs contributing to PLS regression by VIP score, seven were related to *Betaproteobacteria,* five were related to *Anaerolineaceae* (within *Chloroflexi*), and one representative OTU was related to each of *Bellilinea (Chloroflexi), Desulfobacterales (Deltaproteobacteria),* and *Rhizobiales (Alphaproteobacteria).*

**Figure 8.**
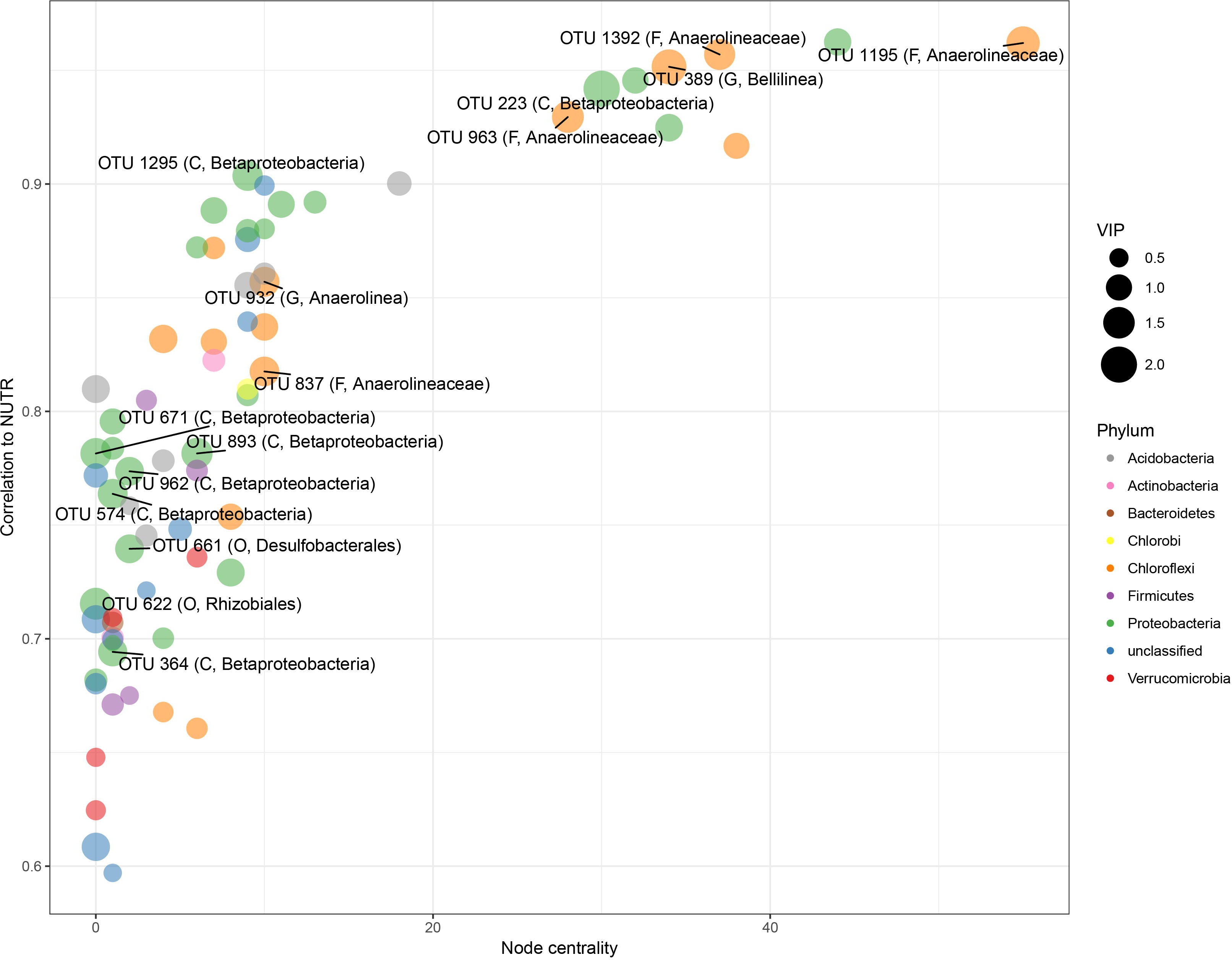
Network visualization and results of partial least squares analysis on the subnetwork most correlated with NUTR. The y-axis represents correlation of OTU to OC values, whereas the x-axis represents the node centrality. Points represent OTUs, and the color of points corresponds to the phylum to which an OTU belongs. Point size corresponds to VIP score of that OTU. The top 15 OTUs are labeled within the graph with corresponding lowest taxonomic identification possible, and the level of that classification. D = Domain; P = Phylum, C = Class, O = Order, F = Family, G = Genus.

For subnetwork relationships to C:N, PLS found that 144 OTUs were 59.0% predictive of variance in C:N (Supplemental Fig. 8). Networks were constructed using an OTU co-correlation threshold of 0.1, within positive or negative correlations (r > +/- 0.2) between OTUs (VIP > 1) and C:N (Fig. 9). Of the top 15 OTUs by VIP score within the network, two OTUs related to *Bacteroidetes* were negatively correlated with C:N. Other top OTUs were positively related to C:N, including seven OTUs related to *Anaerolineaceae,* four OTUs which were unclassified *Bacteria,* and one representative OTU related to each of *Bacillus (Firmicutes*) and *Chloroflexi.*

**Figure 9.**
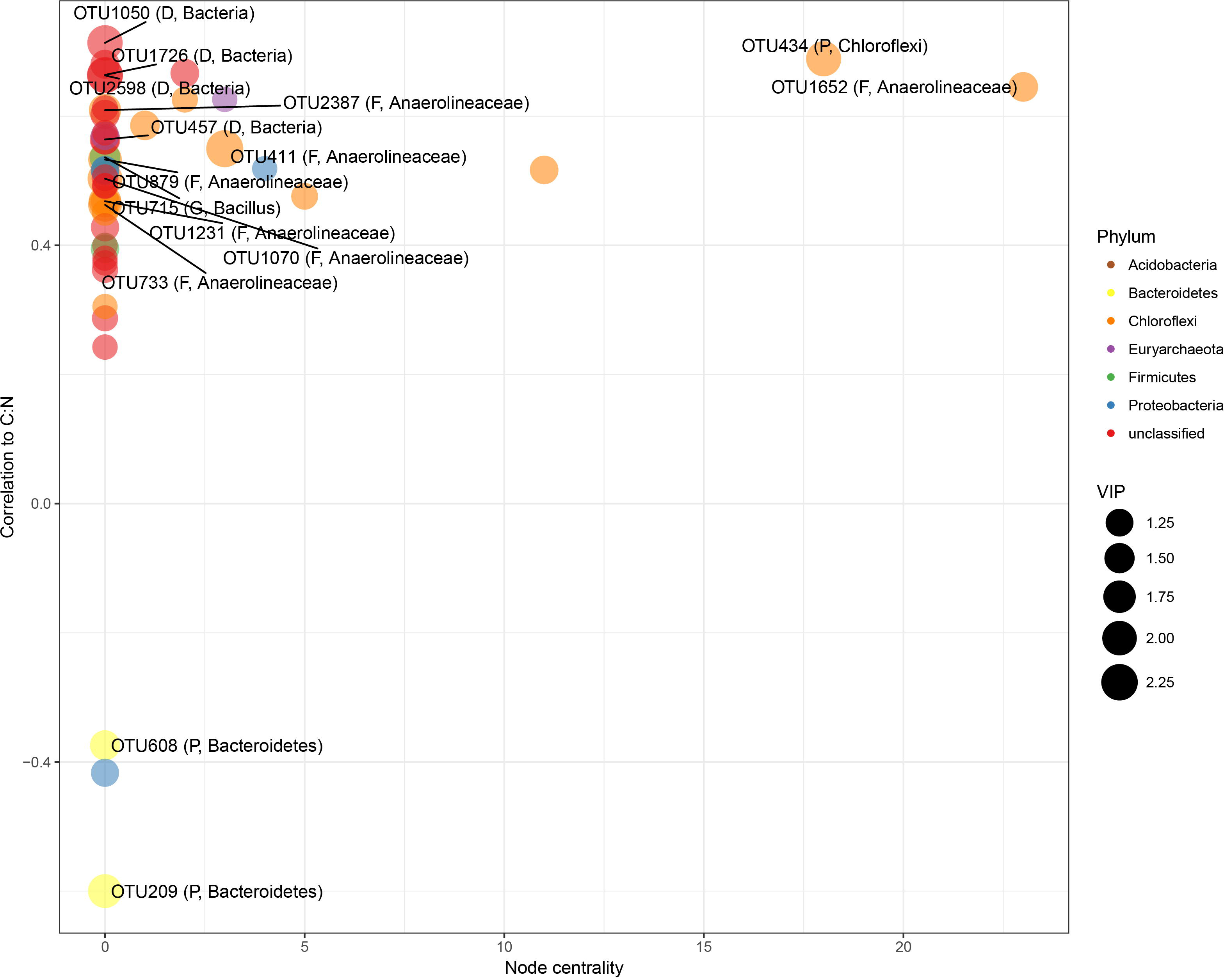
Network visualization and results of partial least squares analysis on the subnetwork most correlated with C:N. The y-axis represents correlation of OTU to C:N, whereas the x-axis represents the node centrality. Points represent OTUs, and the color of points corresponds to the phylum to which an OTU belongs. Point size corresponds to VIP score of that OTU. Only OTUs with a VIP score > 1 were displayed for visualization purposes. The top 15 OTUs are labeled within the graph with corresponding lowest taxonomic identification possible, and the level of that classification. D = Domain; P = Phylum, C = Class, O = Order, F = Family, G = Genus.

## Discussion

### Microbial diversity driven by chemistry within Great Lakes coastal wetlands

This study is the first to suggest that anthropogenic disturbance patterns correspond to microbial community differences in Great Lakes coastal wetlands as consistent with other taxonomic groups such as plants, birds, fish, and invertebrates (Howe *et al.,* 2007; Tulbure *et al.,* 2007; Uzarski *et al.,* 2009; Cooper *et al.,* 2012; Uzarski *et al.,* 2017). Microbial communities appear to respond uniquely to potential anthropogenic influence, as diversity increased with increasing nutrient levels in the coastal wetlands explored in this study. However, microbial community structure was significantly dissimilar between LE and all other wetland regions, and these differences were related to physicochemical differences among coastal wetlands (Fig. 2, Fig. 4, Table 2). As the wetlands within the Lake Erie region maintained the highest nutrient concentrations within the soil, it is possible that anthropogenic stressors related to nutrient loading (and potentially other pollutants) could be driving structural differences in microbial communities among Great Lakes coastal wetlands. Further, network analysis found several taxa/sub communities that were highly correlated to nutrient levels across wetlands explored in this study. Previous research has found that nutrient levels (e.g., C, N, P, etc.), to varying degrees, can influence microbial community composition and structure (Hartman *et al.,* 2008; Peralta *et al.,* 2013; Ligi *et al.,* 2014; Arroyo *et al.,* 2015). Lake Erie coastal wetlands (and the watershed which drains into them) have been historically impacted by anthropogenic pollution and agricultural practices, particularly in comparison to other coastal wetlands within the Laurentian Great Lakes region. This has been demonstrated by multiple ecological indices (e.g., Cvetkovic & Chow-Fraser, 2011; Uzarski *et al.,* 2017) and physicochemical uniqueness (increased levels of nutrients and particulate matter) within the western basin of Lake Erie (Danz *et al.,* 2007; Trebitz *et al.,* 2007; Cvetkovic & Chow-Fraser, 2011; Uzarski *et al.,* 2017). Data presented in this study corroborate this historical evidence of human impact and nutrient loading in the western basin of Lake Erie (Fig. 2., Supplemental Fig. 1), which may be influencing the Lake Erie wetlands explored in this study.

High nutrient influx could also be influencing the chemical and microbial vertical structure within coastal wetland soils. Microbial community and chemical (e.g., C, N, P) vertical structure was not evident within the first 6 cm of soil of coastal wetlands with elevated nutrient levels (e.g. Lake Erie sites). The lack of vertical chemical gradients is unlikely to exclusively explain a corresponding lack of vertical microbial community structure, as some wetland sites lower in nutrient levels also did not experience vertical chemical gradients in this study (e.g. West Saginaw Bay). One possibility is that a lack of vertical chemical structure in conjunction with high nutrient levels in wetland soils could reduce vertical microbial community structure. It has been previously demonstrated that concentrations of carbon electron donors may influence redox gradients within wetland soils (Achtnich *et al.,* 1995), and wetland microbial communities have been demonstrated to correspond with soil redox gradients (Lüdemann *et al.,* 2000; Edlund *et al,* 2008; Lipson *et al,* 2015). However, connections between microbial community metabolic shifts with soil depth and levels of dissolved organic carbon *in situ* remain unresolved in freshwater wetlands (Alewell *et al.,* 2008). Alternatively, another explanation for lack of vertical community structure could be microsite heterogeneity throughout the soil matrix. Previous research in freshwater wetland soils has suggested that microsite heterogeneity may explain coexistence of microbial functional guilds (Alewell *et al.,* 2008; Angle *et al.,* 2017), which could substantially reduce vertical microbial community structural gradients. However, it is necessary to better link microbial community diversity, microbial activity, chemical structure, and microsite heterogeneity to establish relationships between microbial communities and freshwater soil structure. As a caveat, it is possible that chemical and microbial structuring still exists within wetlands with high nutrient levels, yet is not evident within the first 6 cm of soil or at the spatial scale measured in this study. Nevertheless, microbial communities within coastal wetlands with high nutrient levels did not follow the same pattern of vertical structure evident in other comparable coastal wetlands, either chemically or biologically, further suggesting that the integrity of microbial communities within coastal wetland systems may be susceptible to negative anthropogenic pressure.

While relationships between microbial diversity and nutrient levels among coastal wetlands are strong, other unexplored variables unique to Lake Erie (such as geologic history) could also be influencing uniqueness of chemical and microbial profiles in Lake Erie coastal wetlands. The Lake Erie coastal wetland sites explored here were barrier (protected) wetlands, while other wetland sites explored in this study are all classified as lacustrine (open water) wetlands (www.greatlakeswetlands.org). As such, wave action from the Great Lakes impacted wetlands within the western basin of Lake Erie to a lesser degree than other wetlands, thereby reducing sediment export rates into the Great Lakes themselves. Hydrologic energy was found to impact wetland primary productivity and respiration in Lake Huron coastal wetlands, suggesting Great Lakes ecosystems may exert unique environmental forces on wetland microbial communities (Cooper *et al.,* 2013). Low carbon export rates or elevated sedimentation rates may exist in the western basin of Lake Erie as consequence of low wave action in these wetlands, which may influence the chemical and biological structure (such as vertical microbial community structure) within wetland soils of this region. Nevertheless, previous research at the same wetland locations explored in this study have demonstrated that wetlands within the western basin of Lake Erie are highly degraded with respect to other wetlands (Uzarski *et al.,* 2017), particularly with respect to physicochemical conditions. Additionally, the same vegetation zone (dominated by cattails or bulrush) was sampled among all wetlands explored in this study as an attempt to reduce bias in distinct environmental conditions which may exist in other vegetation zones among wetland sites. Burton et al. (2002) suggested that soil organic content was related to plant zonation in Great Lakes coastal wetlands. Further research would be necessary to fully tease apart the effects of anthropogenic stress and other natural contributions to differences in microbial communities among coastal wetlands.

### Taxonomic patterns among wetland regions and soil depths

At the level of phylum, Great Lakes coastal wetlands shared many similarities regardless of environmental conditions (Supplemental Fig. 4), and shared dominant groups such as *Deltaproteobacteria, Betaproteobacteria, Gammaproteobacteria, Bacteroidetes,* and *Chloroflexi*. These bacterial groups have been commonly found within other wetland soils (Hartman et al., 2008; Ansola et al., 2014; Arroyo et al., 2015). However, there were distinct differences in community composition among wetland regions as demonstrated by perMANOVA and NMDS, particularly between LE and all other regions. More specifically, several *Planctomycetes* OTUs were less abundant within LE than within other wetland regions (Fig. 6), suggesting this taxonomic group may thrive in less impacted wetland soils. This pattern was similar for other groups of bacterial taxa, including *Spartobacteria, Sphingobacteria, Clostridia,* and *Caldilineae,* as well as archaeal taxa including methanogenic *Methanomicrobia* such as *Methanocella, Methanoregula, Methanosaeta* and *Methanosarcina.* It is important to recognize that, while unique patterns in archaeal diversity were found among wetland regions, primers employed in this study were not designed to explore archaeal diversity, and thus this representation of archaeal diversity is likely incomplete. Several Acidobacteria OTUs were uniquely abundant in LE wetlands (e.g. Acidobacteria Groups 6, 17, and 18). Acidobacterial abundance has been shown to increase with decreasing pH within soil environments (Jones et al., 2009), and as such, the relatively lower pH of LE soils with respect to other wetland regions may be driving this trend within freshwater coastal wetlands.

Several other taxonomic groups of microbes were differentially abundant among wetland soil habitats, often dependent on soil depth. Perhaps most interestingly, archaeal OTUs within *Chrenarchaeota* and *Euryarchaeota* were more relatively abundant in soils between 4 – 6 cm in depth, particularly within Classes *Thermoprotei, Methanomicrobia,* and *Methanobacteria.* Many of these OTUs were identified to the genus level, including the methanogenic *Methanosaeta*, *Methanoregula*, and *Methanobacterium*. Recent research has suggested that methanogenic activity can often be highest within oxygenated soils, which can occur within the top 10 cm of freshwater wetland soils (Angle et al., 2017). As soils within our study were sampled to a maximum depth of 6 cm, it is possible that methanogens within Great Lakes coastal wetlands may be active in the oxygenated layer of soils, particularly between 4 – 6 cm where oxygen, while possibly present, is lower than layers of soil directly above. However, oxygen was not measured within the soil of this study, and thus further research would be necessary to understand whether oxygen is permeating to 4 cm depth in wetland soils explored herein. Within the top 0 – 2 cm of soil, several bacterial OTUs were differentially abundant, most notably within taxonomic groups such as *Alpha-, Beta-,* and *Gammaproteobacteria,* several groups of *Acidobacteria, Spartobacteria, Verrucomicrobiae, Planctomycetes,* and *Sphingobacteria.*

### Relationships between microbial subnetworks and environmental gradients

Through network analyses, multiple subcommunities were delineated which were significantly related to environmental gradients (such as nutrients C, N, and P) among coastal wetlands sampled in this study. Specifically, a subnetwork of 69 microbial taxa was 93.8% predictive of nutrient level variation among coastal wetland soils. Several microbial taxa within this subcommunity were individually predictive of nutrient levels to a high degree, including several OTUs within *Anaerolineaceae,* one OTU within genus *Anaerolinea,* and another within genus *Bellilinea.* From the genus *Anaerolinea,* two thermophilic chemoorganotrophs *(Anaerolinea thermophila* and *Anaerolinea thermolimosa*) have been isolated (Sekiguchi *et al.,* 2003; Yamada *et al.,* 2006). Only one isolated member has been established within the genus *Bellilinea (Bellilinea caldifistulae*); it has been described as a thermophilic, fermentative, obligate anaerobe which thrives in co-culture with methanogens (Yamada *et al.,* 2007). It is unlikely that the OTUs found in our study are the same species as the isolated *Anaerolinea* and *Bellilinea* species, as coastal wetland soils are not high-temperature environments necessary for thermophilic species. Additionally, no OTUs related to methanogenic archaea were found within this subnetwork, suggesting that *Anaerolineacea* OTUs within coastal wetland soils may fluctuate independently of any specific methanogenic OTUs. It is possible that the *Bellilinea* OTU found within the subnetwork is related to nutrient level concentrations. This would support fermentative metabolism as noted within *Bellilinea caldifistulae.* It is important to note that several other studies have discovered OTUs related to *Anaerolineaceae* within wetland soils, with upwards of 90% relative abundance among *Chloroflexi* OTUs within these systems (Ansola *et al.,* 2014; Deng *et al.,* 2014; Hu *et al.,* 2016). This suggests that there are probable mesophilic species yet to be isolated within this ubiquitous family of bacteria, which may be of high importance within wetland soils. Interestingly, the majority of OTUs (61 out of 69 OTUs) within the subnetwork most related to NUTR shifts were also differentially abundant between LE and all other regions (Fig. 6). The parallels drawn between these two analyses highlights the potential importance of NUTR (NH_4_+, OM, OC, TN) in driving differences in microbial OTU abundances between LE and other coastal wetland regions.

*Betaproteobacteria* were also found to significantly predict nutrient levels among coastal wetlands. Hu *et al.* (2016) found that both *Betaproteobacteria* and *Anaerolineae* were positively related to TN levels, which is consistent with the data presented here, and these two taxa were suggested to contribute to higher levels of heterotrophic activity. Further, *Anaerolineaceae* OTUs were consistently related to increasing C:N, suggesting that many taxa within this family have preference for recalcitrant carbon sources. As C:N also tends to increase with soil depth, it is also probable that the putatively obligate anaerobic *Anaerolineaceae* are coinciding with decreasing oxygen levels and/or changing metabolism requirements with increasing soil depth.

Development of biological indices and establishment of indicator taxa have been suggested as necessary for microbial communities within wetlands (Uzarski *et al.,* 2017), particularly through the use of high-throughput sequencing technologies which now allow for deep assessment of microbial community composition and structure within environmental samples (Sims *et al.,* 2013; Urakawa & Bernhard, 2017). Specifically within Great Lakes coastal wetlands, it is integral to develop ecosystem health indicators based upon multiple different groups of taxonomy, as separate biological indices can present contrasting assessments of wetland health (Uzarski *et al.,* 2017). As microbial indicators have yet to be established in Great Lakes coastal wetlands, this research begins the first steps in exploring how microbial communities can be used as an additional and potentially important ecosystem health indicator. In addition to their importance as biological signals for environmental health, microbial indicator taxa may play prominent roles in bioremediation of excess nutrients and pollutants found within anthropogenically impacted coastal wetlands. Network analyses in this study have allowed for the generation of hypothetical subcommunities of diverse microbial taxa related to nutrient levels among Great Lakes coastal wetlands, and could assist in further understanding of which microbial taxa may be responding to anthropogenic stress in these ecosystems.

## Conclusions

This study marks the first characterization of microbial communities within Great Lakes coastal wetlands. Coastal wetlands are integral in the proper functioning of biogeochemical cycles and environmental sustainability of the Great Lakes. While it has long been known that anthropogenic pressure can impact animal and plant communities within these coastal wetlands, this is the first evidence that these pressures may also be influencing microbial communities and may be influencing biogeochemical cycles by extension. Alpha and beta diversity were both related to nutrient gradients among and within regions, suggesting that variability in microbial community structure is highly coupled to geochemistry within wetland soils. We propose that wetland microbial community structure can also potentially be used to assess a wetland for monitoring purposes. As illustrated within this study, wetland microbial community structure and depth are decoupled within the wetlands experiencing the highest nutrient levels, likely originating from terrestrial inputs due to human activity. As such, multivariate statistics (as used in the methods of this study) may prove useful in examining relationships between wetland soil depth and microbial community structure alongside microbial network analyses, which could provide biological indicators of nutrient loading stress on coastal wetland habitats. We propose that wetland microbial community structure can also potentially be used to assess a wetland for monitoring purposes.

Further, this study provides insight on microbial community subnetworks and individual OTUs, which were predictive of chemical concentrations, and may be useful for future management of Great Lakes coastal wetland systems. Within subnetworks existed multiple taxa with strong individual relationships to environmental gradients among coastal wetlands throughout the Great Lakes. Even further, several community members within these subnetworks were taxonomically related (such as OTUs related to *Anaerolineaceae* within *Chloroflexi*), suggesting that specific taxonomic groups of microbes may be useful to explore further as potential biological indicator groups. This study highlights the strength of network analyses (such as WGCNA) in delineating hypothetical networks of interacting microbes, and whether these networks are predictive of physical or chemical gradients measured within an environment.

## Acknowledgements

Special thanks to Alexandra Bozimowski and Thomas Langer for their assistance in collecting the wetland samples and data acquisition, as well as Dr. Matthew Cooper for providing valuable insight and conversation on wetland ecology and the custom-built core extruder built by Mr. Gary Cooper. Special thanks also to Mike Henson for providing assistance on network statistical analyses. This paper is Contribution Number XX of the Central Michigan University Institute for Great Lakes Research.

